# Connective-Tissue Growth Factor (CTGF/CCN2) Contributes to TGF-β1-Induced Lung Fibrosis

**DOI:** 10.1101/2020.07.04.187492

**Authors:** Toyoshi Yanagihara, Sy Giin Chong, Mahsa Gholiof, Kenneth E. Lipson, Quan Zhou, Ciaran Scallan, Chandak Upagupta, Jussi Tikkanen, Shaf Keshavjee, Kjetil Ask, Martin RJ Kolb

## Abstract

Idiopathic pulmonary fibrosis (IPF) is a fatal lung disease characterized by progressive and excessive accumulation of myofibroblasts and extracellular matrix in the lung. Connective-tissue growth factor (CTGF) is known to exacerbate pulmonary fibrosis in radiation-induced lung fibrosis, and in this study, we show the upregulation of CTGF from a rat lung fibrosis model induced by adenovirus vector encoding active TGF-β1 (AdTGF-β1), and also in patients with IPF. The expression of CTGF was upregulated in vascular smooth muscle cells cultured from fibrotic lungs on days 7 or 14 as well as endothelial cells sorted from fibrotic lungs on day 14 or 28 respectively. These findings suggest the role of different cells in maintaining the fibrotic phenotype during fibrogenesis. Treatment of fibroblasts with recombinant CTGF along with TGF-β increases pro-fibrotic markers in fibroblasts, confirming the synergistic effect of recombinant CTGF with TGF-β in inducing pulmonary fibrosis. Also, fibrotic extracellular matrix upregulated the expression of CTGF, as compared to normal extracellular matrix, suggesting that not only profibrotic mediators but also a profibrotic environment contributes to fibrogenesis. We also showed that pamrevlumab, a CTGF inhibitory antibody, partially attenuates fibrosis in the model. These results suggest that pamrevlumab could be an option for the treatment of pulmonary fibrosis.

## Introduction

Idiopathic pulmonary fibrosis (IPF) is a fatal progressive disease with an unknown etiology that is characterized by excessive accumulation of myofibroblasts and extracellular matrix in the lungs, and with a median survival of 3–5 years after diagnosis (1)(2). Currently, there are two available FDA-approved drugs for IPF: nintedanib and pirfenidone (3). These drugs are thought to inhibit pathways in the aberrant wound healing response and to slow disease progression, but fail to stop or reverse IPF progression. We therefore need to elucidate key pathways in IPF progression to effectively target and treat this fibrotic disease.

Connective-tissue growth factor (CTGF), also known as CCN2 (Cyr61 [cysteine-rich protein 61], CTGF [connective tissue growth factor] and NOV [nephroblastoma overexpressed gene]; and more recently coined, cellular communication network 2), is a secreted protein with a role in different cellular events, such as cell proliferation, differentiation, migration, skeletogenesis, angiogenesis, and wound healing (4)(5). CTGF expression is regulated by various growth factors and cytokines, including transforming growth factor (TGF-β) and bone morphogenetic protein (BMP). CTGF has a cysteine knot domain that mediates its interaction with cell surface and matrix heparan sulfate proteoglycans, and keeps it in the vicinity of the cells from which it is secreted, enabling it to act in an autocrine or paracrine manner. It has been suggested that CTGF plays a role in extracellular matrix (ECM) production due to its ability for mediating collagen deposition in wound healing. CTGF is also involved in many pathological conditions. CTGF leads to myofibroblast formation by transdifferentiating other cells, such as resident fibroblasts, and epithelial cells. CTGF also has a role in remodeling and deposition of ECM by activating myofibroblasts which leads to fibrosis and tissue remodelling. When vasculature undergoes tissue remodeling, it can lead to local hypertension which itself increases CTGF expression. This creates a positive feedback loop that leads to more tissue remodeling. CTGF also increases the expression of different cytokines, including VEGF and TGF-β? which in turn increase CTGF expression, creating more positive feedback loops as a result.

In this study, we showed upregulation of CTGF in lung tissues from patients with IPF, as well as in a rat lung fibrosis model induced by adenoviral vector encoding TGF-β1^223/225^ (originally named AdTGF-β1^223/225^; hereafter called AdTGF-β1) harboring a mutation of cysteine to serine at positions 223 and 225, with the expressed TGF-β1 biologically active (6). This model has been shown to induce more intense, rapid and severe fibrotic reactions, with greater increases in hydroxyproline, than those reported in various bleomycin studies (7). Lung fibrosis begins on day 7 and persists until day 28. Less pronounced acute lung injury, particularly to the epithelium, and accompanying inflammation were also evident in this model, compared to the bleomycin model (7). We also showed that pamrevlumab, a CTGF inhibitory antibody, partially attenuates fibrosis in the model.

## Materials and Methods

### Ethics approval and consent to participate

All animal work was conducted under the guidelines from the Canadian Council on Animal Care and approved by the Animal Research Ethics Board of McMaster University under protocol #17-07-31. All procedures using human tissues were approved by the Hamilton Integrated Research Ethics Board (11-3559, 13-523-C).

### Human samples

Control lung tissues were from patients undergoing a surgical procedure for cancer. IPF lungs were from patients undergoing explant at Toronto Lung Transplant Programme. The lung samples were about 1–2 cm in size.

### Animal Experiments

Pulmonary fibrosis was induced by an adenoviral gene vector encoding biologically active TGF-β1 (AdTGF-β1). Female Sprague-Dawley rats (250–300 g; Charles River, Wilmington, MA) received 5.0 × 10^8^ PFU of AdTGF-β1 by single intratracheal instillation under isoflurane anesthesia on day 0. Control rats received an empty vector construct (AdDL). Pamrevlumab (FibroGen, 25 mg/kg) or isotype control antibody (FibroGen, 25 mg/kg) was given intraperitoneally from day 10 to day 27 three days in a week in a double-blinded manner. Rats were sacrificed on day 7, 14, or 28 and lung tissue was harvested.

### Antibodies and reagents

Antibodies used were anti-α-smooth muscle actin (α-SMA) (abcam, ab7817), anti-α-tubulin (Cell signaling technology, #2144S), anti-CTGF (Santa Cruz, sc-365970), anti-Collagen type I (Rockland, #600-401-103-0.1), anti-TGF-β (Cell signaling technology, #3709S), anti-Smad3 (Cell signaling technology, 9513S), anti-phospho-Smad3 (pSmad3) (Cell signaling technology, #9520S), anti-Sm22 (abcam, ab14106), anti-Vimentin (abcam, ab16700), anti-VWF (LS Bio, LS-C411685-100), anti-rabbit HRP linked IgG (Cell Signaling Technology, #7074), Anti-mouse IgG HRP-linked Antibody (Cell Signaling Technology, #7076). For fluorescence microscopy, we used goat or donkey secondary antibody conjugated with Alexa Fluor-488 and Alexa Fluor-594 (abcam) as secondary antibodies. Recombinant human CTGF was from FibroGen. Recombinant human TGF-β1 (#240-B) was purchased from R&D systems.

### Cell preparation and culture

Human normal lung fibroblasts (HLFs) (PCS-201-013) were purchased from ATCC and cultured in 10% Fetal Bovine Serum (FBS) and 1% pen-strep (Gibco, Life Technologies) in Dulbecco’s modified Eagle Medium (DMEM) (Biowhittaker® Reagents, Lonza). For primary rat fibroblasts, single-cell suspensions were created by mincing and 1 hr digestion with 1 mg/ml of collagenase type I (Sigma-Aldrich, C0130-1G) and 20 µg/ml of DNase I (Sigma-Aldrich, 11284932001) of harvested rat lungs, then cells were cultured for 7 days or till confluency and used as primary fibroblasts. For primary rat vascular smooth muscle cells (VSMCs), central pulmonary arteries from AdDL or AdTGF-β1-treated lungs were carefully dissected and cut into < 1 mm pieces and cultured in 10% FBS and 1% pen-strep in DMEM in 6 well plates for 8 days. VSMCs were characterized by being positive for Sm22 (>95%) and negative for endothelial cell marker CD31 by immuno-staining. HLFs were used at passages between P2 and P6. Primary rat VSMCs were used at passage P1. All cells were incubated at 37°C, 5% CO_2_.

### Histology

The left lobes of rat lungs were fixed by intratracheal instillation of 10% neutral-buffered formalin at a pressure of 20 cm H_2_O. Picture acquisition of Masson’s Trichrome-stained (MT-stained) sections was performed using an automatic slide scanner microscope (Olympus VS 120-L).

### Ashcroft score

Pulmonary fibrosis of Masson’s Trichrome-stained lung sections was graded from 0 (normal lung) to 8 (completely fibrotic lung), using a modified Ashcroft score (8).

### Immunofluorescence

Immunostaining was performed on formalin-fixed rat lung tissue sections. Briefly, following deparaffinization and saturation of nonspecific sites with 10% horse serum/PBS for 30 min, lung sections were incubated with primary antibodies overnight in a humidified chamber at 4°C. Conjugated secondary antibodies were used at a dilution of 1:500. Slides were mounted in Prolong-gold with DAPI (ProLong® Gold antifade reagent with DAPI, Life technologies, P36931).

### Western blotting

Crushed lungs were homogenized using a mechanical homogenizer (Omni International, Waterbuy CT) or cultured cells were lysed in 1× lysis buffer (NEB, #9803S) supplemented with complete protease inhibitors (Roche) and the collected supernatant was used for western blotting. Total proteins from lung homogenate (30–60 µg) or cells (20 µg) were separated on a 6–12% SDS Polyacrylamide Electrophoresis gels based on the molecular weight. Proteins were transferred to a PVDF membrane (Bio-Rad Laboratories) using a wet transfer apparatus and blocked at room temperature for 30 min using 5% skim milk. Protein detection was performed using Clarity(tm) Western ECL Substrate (Bio-Rad Laboratories) and read in a ChemiDoc XRS Imaging System (Bio-Rad Laboratories). The signals were measured using ImageJ public-domain software.

### Flow cytometry sorting

Enzymatically digested rat lung was treated with ACK lysis buffer to eliminate erythrocytes and washed with PBS, followed by incubation with anti-Fcγ III/II receptor. Anti-rat CD31-viobright FITC (Miltenyi, 130-105-880), anti-rat CD45-APC/Cy7 (BD Biosciences, 561586), and 7-Aminoactinomycin D (7-AAD) (BD, 559525) were used. Flow cytometry sorting was done on MoFlo XDP (Beckman Coulter).

### Isolation of mRNA and gene expression

Total RNA was extracted from frozen lung tissue or cultured cells with TRIzol® reagent (Thermo fisher scientific, 15596026) according to the manufacturer’s recommendations. For sorted ECs, total RNA was extracted using RNeasy Plus Micro Kit (Qiagen, 74034). Total RNA was reverse transcribed using qScript cDNA SuperMix (Quanta Bioscience, 95048-025, Gaithersburg, MD). The cDNA was amplified using a Fast 7500 real-time PCR system (AB Applied Biosystems) using TaqMan® Universal PCR Master Mix and predesigned primer pairs (Thermo Fisher Scientific) for rat *Gapdh* (Rn01775763_g1), rat *Ctgf* (Rn01537279_g1), human *ACTA2* (Hs00426835_g1), human *COL1A1* (Hs00164004_m1), human *CTGF* (Hs00170014_m1), human *TGFBR1* (Hs00610320_m1), and human *GAPDH* (Hs02786624_g1).

### Decellularization and Recellularization of Rat Lungs

Lungs were harvested on day 28 after AdTGF-β1 or AdDL administration. Decellularization was performed through manual lung perfusion (10 ml each time) and incubation with 50 ml of: TritonX (1%; overnight), Sodium Deoxycholate (2%; overnight), NaCl (1 M; 1 hr), and stored in PBS supplemented with Penicillin Streptomycin (1%; max 3 months). Lungs were washed with sterile H_2_O between each solution during decellularization to ensure there was no mixing of wash solutions. Left lobe recellularization was performed by first tying off the right lobes leaving only the left lobe open. The left lobe was perfused with 5 ml of fibroblasts (1×10^6^ cells/ml) in 2% low melting agarose. Lungs were briefly left to solidify at room temperature and the left lobe was sectioned into horizontal slices of ∼3 cm and immersed in 10% FBS and 1% pen-strep in DMEM and incubated at 37°C, 5% CO_2_ for 1 and 7 days.

### Data mining of single-cell RNA sequencing on human lungs

Scatter plot with bar of CTGF expression across different cells in human lungs from healthy subjects (n=29) and patients with IPF (n=32) by single cell RNA sequencing. The data were generated from http://ipfcellatlas.com/ (9).

### Statistical Analysis

The Student’s two-tailed unpaired t-test was used to compare two groups. Statistical analysis between multiple groups with one control group was performed by one-way analysis of variance (ANOVA) followed by Dunnett’s multiple comparison test with the use of GraphPad Prism 8 (GraphPad Software Inc.). A *p*-value of less than 0.05 was considered significant.

## Results

### CTGF was upregulated in lungs from patients with IPF

The expression of CTGF in lungs from patients with IPF was first examined by immunofluorescence. As expected, CTGF expression was elevated in the IPF lung and co-localized with Vimentin positive cells (fibroblasts) and VWF positive cells (ECs) (Figure 1A). Western blots of explanted lung lysates showed a tendency for increased CTGF expression in patients with IPF compared to control lungs (1.43±0.75 [mean ± SD], *p* = 0.09), that did not achieve statistical significance due to the high variability in the IPF lungs (Figure 1B, 1C). We classified IPF patients into subgroups: IPF + no treatment, IPF + pirfenidone, and IPF + nintedanib. Treatment with pirfenidone or nintedanib had a tendency to suppress CTGF expression in IPF lungs compared to IPF without treatment, again though not significant (IPF: 1.62±0.76, IPF + pirfenidone: 1.38±0.83, IPF + nintedanib: 1.36±0.56. [mean ± SD])(Figure 1D).

**Figure 1.**
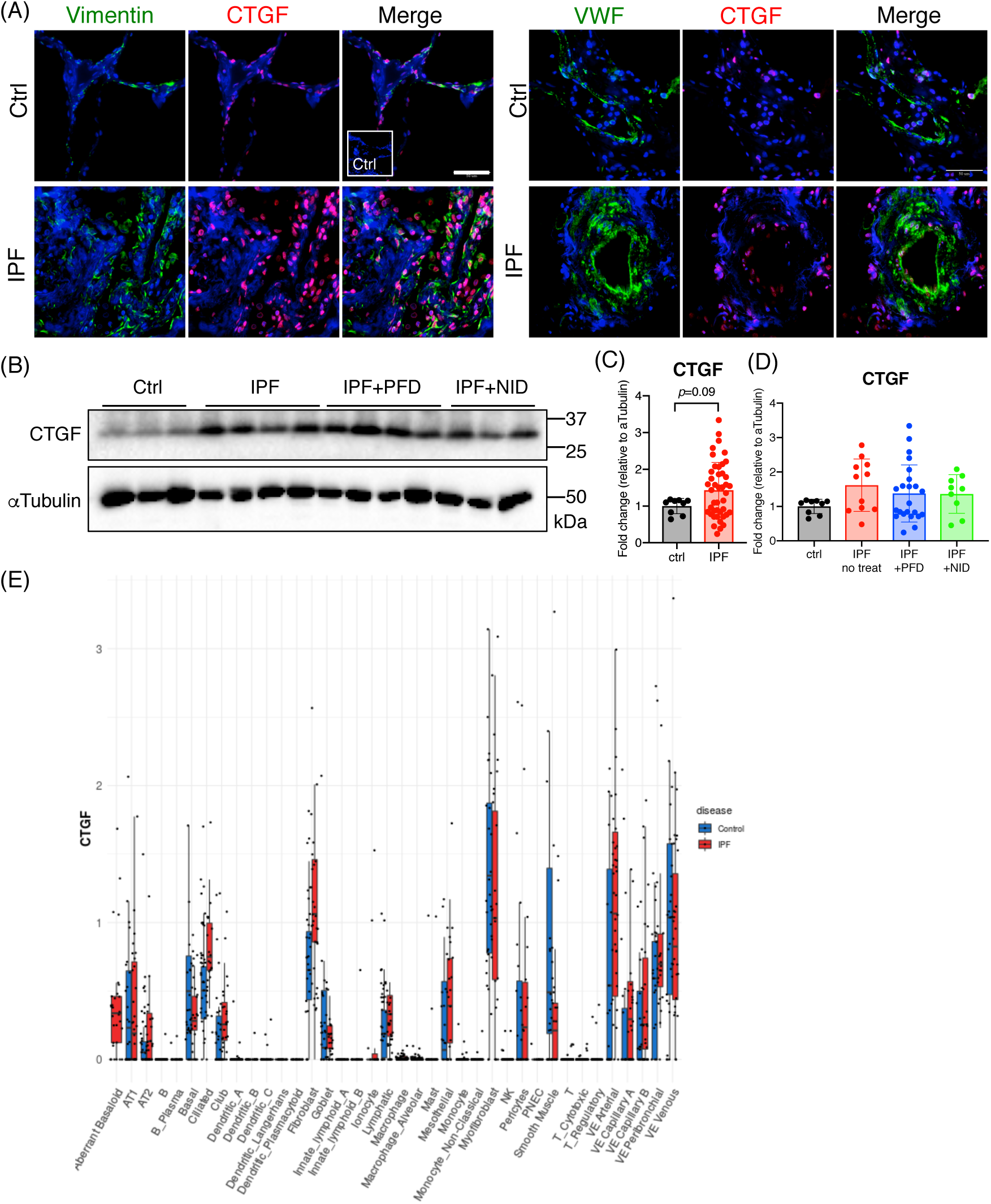
CTGF expression in the human lung. (A) Immunofluorescence staining of Vimentin, VWF, CTGF, and DAPI on the lung sections from a control or a patient with idiopathic pulmonary fibrosis (IPF). (B) Western blot analysis and quantification (C) for CTGF and αTubulin in explanted lung lysates of patients with IPF. (D) IPF patients were sub-grouped into IPF+no treatment, IPF+PFD, and IPF+NID. Ctrl: n=9, IPF+no treatment: n=11, IPF+PFD: n=23, IPF+NID: n=9. PFD; pirfenidone, NID; nintedanib. Data are expressed as mean ± SD. The amount of CTGF in untreated IPF lungs were compared to that of control, IPF+PFD, and IPF+NID by ANOVA with Dunnet’s multiple comparison test. (E) Scatter plot with bar of CTGF expression across different cells in human lungs from healthy subjects (n=29) and patients with IPF (n=32) by single cell RNA sequencing. The data were generated from http://ipfcellatlas.com/.

Recent advances in single-cell RNA sequencing (scRNA-seq) have unveiled cell-specific gene expression in the lung from healthy subjects (n=14) and patients with IPF (n=26) (9). Mining of the scRNA-seq data showed *CTGF* was expressed in several cell types, including fibroblasts, myofibroblasts, smooth muscle cells, vascular endothelial cells, ciliated cells, basal cells, type 1 alveolar epithelial cells, mesothelial cells, and lymphatic cells (Figure 1E).

### CTGF was upregulated in AdTGF-β1-induced lung fibrosis in rats

To investigate the association CTGF with pulmonary fibrosis in a rodent model, female Sprague-Dawley rats were exposed to AdTGF-β1 by a single intratracheal instillation on day 0. Control rats received an empty vector construct (AdDL). Immunofluorescence (IF) staining of lung sections showed that expression of CTGF was upregulated in the AdTGF-β1 rats, as shown in Figure 2A. This result was confirmed with Western Blot analysis for CTGF in whole lung lysates of AdDL and AdTGF-β1 rats (Figure 2B, 2C). It is well known that fibroblasts express CTGF (10), which was also supported by scRNA-seq data from human lungs (Figure 1C). In addition to fibroblasts, vascular cells also appeared to express CTGF, including endothelial cells and Sm22-positive vascular smooth muscle cells (Figure 2A).

**Figure 2.**
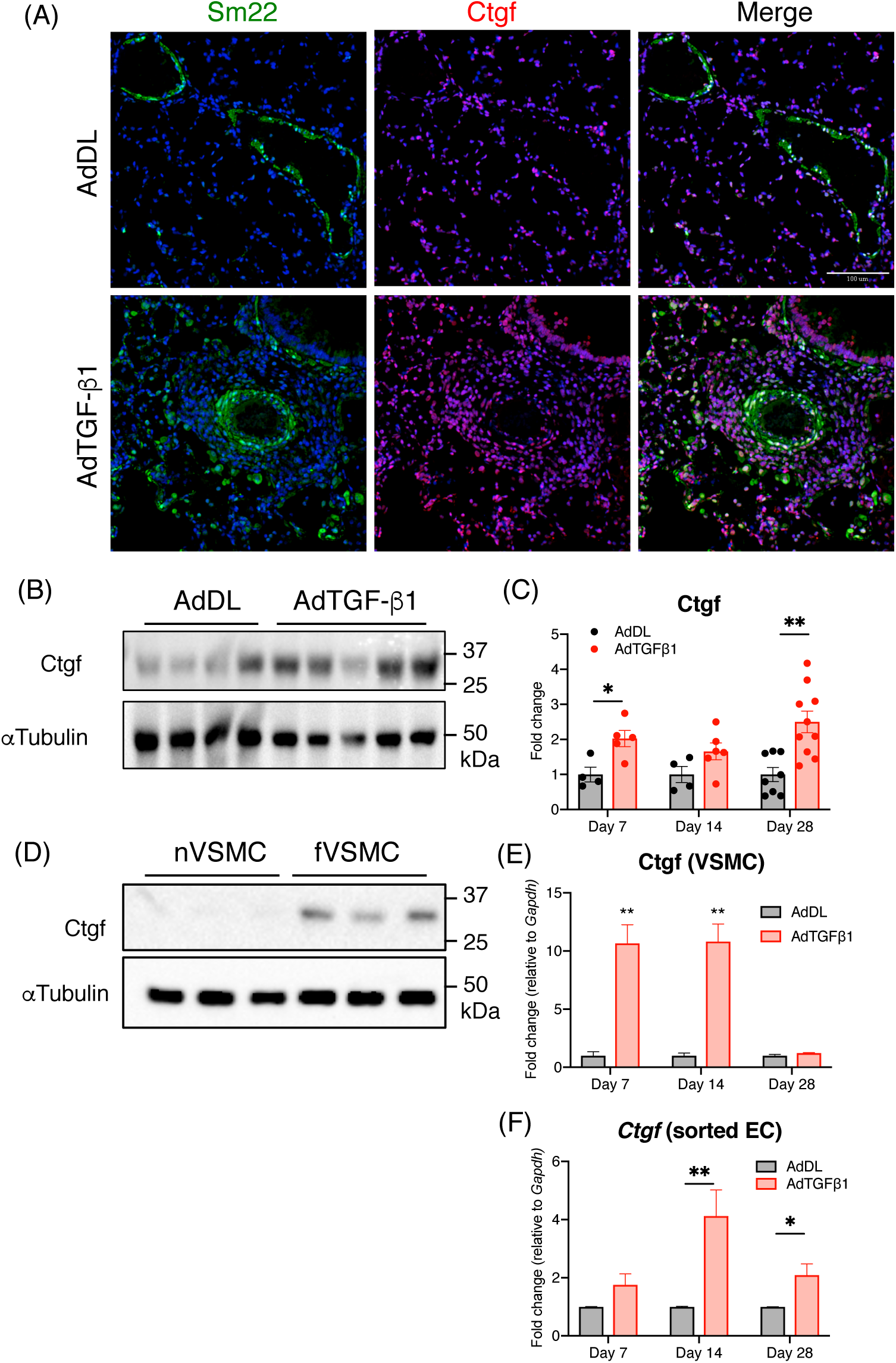
CTGF was upregulated in AdTGF-β1-induced lung fibrosis in rats. (A) Immunofluorescence staining of Sm22, CTGF, and DAPI on the lung sections from rats treated with AdDL and AdTGF-β1 on day 7. (B) Western blot analysis and quantification (C) for CTGF and αTubulin in whole lung lysates of AdDL and AdTGF-β1. Representative images are on day 7. (D) Western blot analysis and quantification (E) for CTGF and αTubulin in the cultured vascular smooth muscle cells (VSMCs) from AdDL (nVSMCs) and AdTGF-β1 (fVSMCs) on days 7, 14, and 28 (*n*=3). (F) mRNA expression of *Ctgf* in the sorted endothelial cells (ECs) from AdDL or AdTGF-β1-treated rat lungs (*n*=4 for each group). Data are expressed as mean ± SEM. \*\**p* < 0.01, \**p* < 0.05.

VSMCs were isolated from the pulmonary arteries in lungs of AdDL (nVSMCs) and AdTGF-β1 (fVSMCs) rats on 7, 14 and 28 days after infection and cultured. CTGF expression was upregulated in fVSMCs isolated from lungs on days 7 or 14 but not on day 28 (Figure 2D, 2E), suggesting VSMCs contribute to CTGF expression in the early stages of fibrosis. On the other hand, increased CTGF expression was observed in sorted endothelial cells (ECs) isolated from AdTGF-β1 rats on days 14 and 28, suggesting ECs contribute to CTGF expression in the middle to late stages of fibrosis (Figure 2F).

### Recombinant CTGF synergistically induced pro-fibrotic markers in fibroblasts along with TGF-β

Next, we sought to examine the pro-fibrotic effect of CTGF on fibroblasts. Human lung fibroblasts were treated with recombinant CTGF for 24 hours under serum free condition. The mRNA expression of pro-fibrotic markers such as *ACTA2* and *COL1A1* were examined. Although the mRNA expression of *ACTA2* and *COL1A1* were shown to be slightly increased as they were treated with increasing concentration of recombinant CTGF (Figure 3A), high concentrations of recombinant CTGF were required (over 1∼3 µg/ml) to observe significant effects. To determine whether recombinant CTGF works synergistically with TGF-β, expression of the profibrotic markers was also examined after the human lung fibroblasts were treated with recombinant TGF-β with and without recombinant CTGF for 24 hours. The expression of *ACTA2, COL1A1, and CTGF* was greatly elevated when human lung fibroblasts were treated with recombinant TGF-β in combination with low concentrations of recombinant CTGF that induced no *ACTA2* expression on their own (Figure 3B). This observation indicates a synergistic effect between these two factors and suggests a substantial role for CTGF in mediating fibrosis.

**Figure 3.**
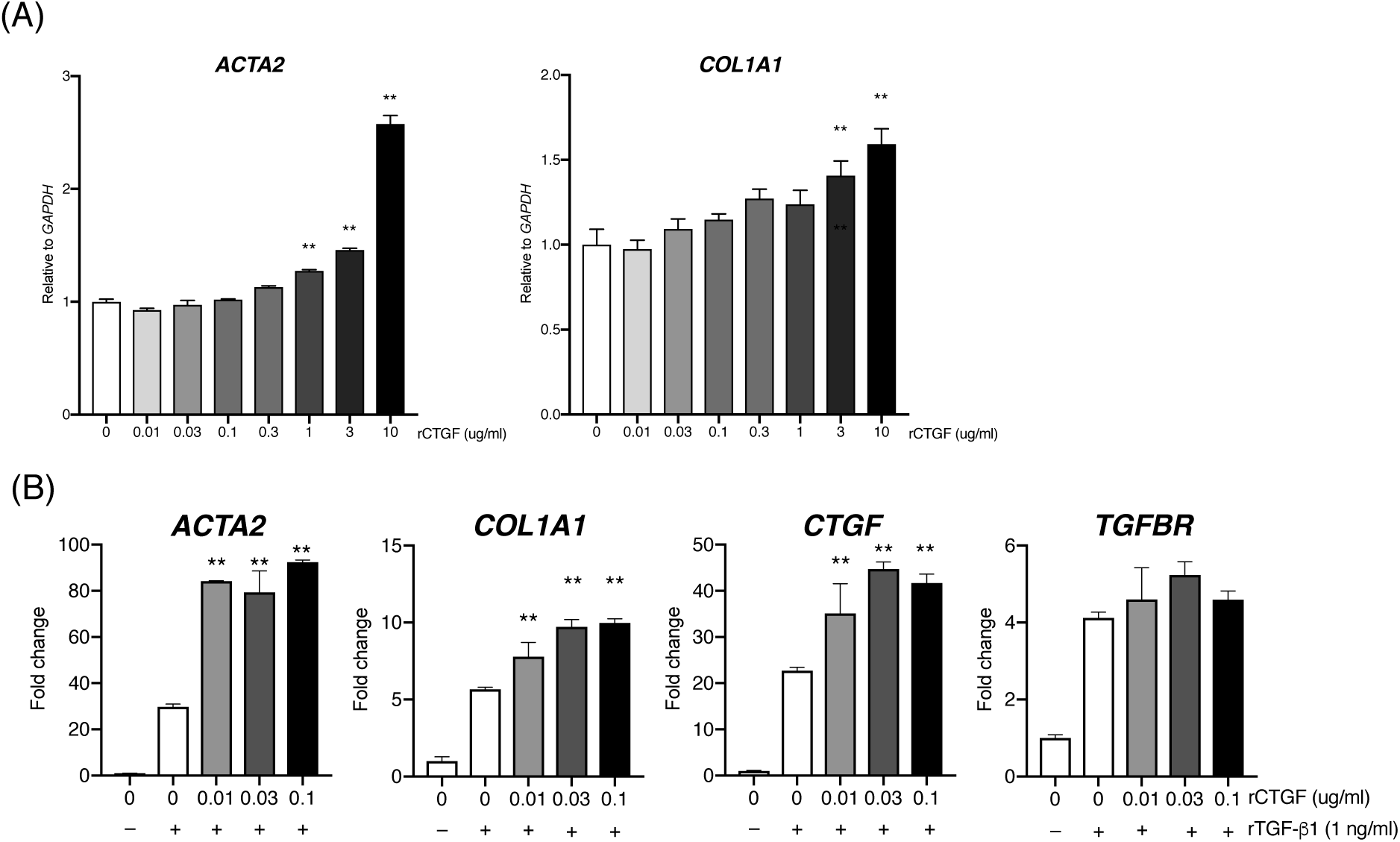
Recombinant CTGF synergistically induced pro-fibrotic markers in fibroblasts along with TGF-β. (A) mRNA expression of *ACTA2* and *COL1A1* in human lung fibroblasts treated with recombinant CTGF for 24 hrs under serum free condition. Levels of gene expression were compared with that in fibroblasts without recombinant CTGF treatment. (B) mRNA expression of *ACTA2, COL1A1, CTGF, TGFBR* in human fibroblasts treated with recombinant TGF-β with/without recombinant CTGF for 24 hrs. Levels of gene expression were compared with that in fibroblasts treated with recombinant TGF-β Without recombinant CTGF. \*\**p* < 0.01, \**p* < 0.05.

### The regulatory mechanism of CTGF in the cells

To explore regulatory mechanisms of CTGF expression in the fibroblasts, cells were cultured under different conditions *in vitro* and *ex vivo*. To determine the TGF-β dose-response, human lung fibroblasts were stimulated with different concentrations of recombinant TGF-β for 24 hours under serum free condition. The expression of *CTGF* mRNA increased with increasing concentrations of TGF-β stimulation (Figure 4A).

**Figure 4.**
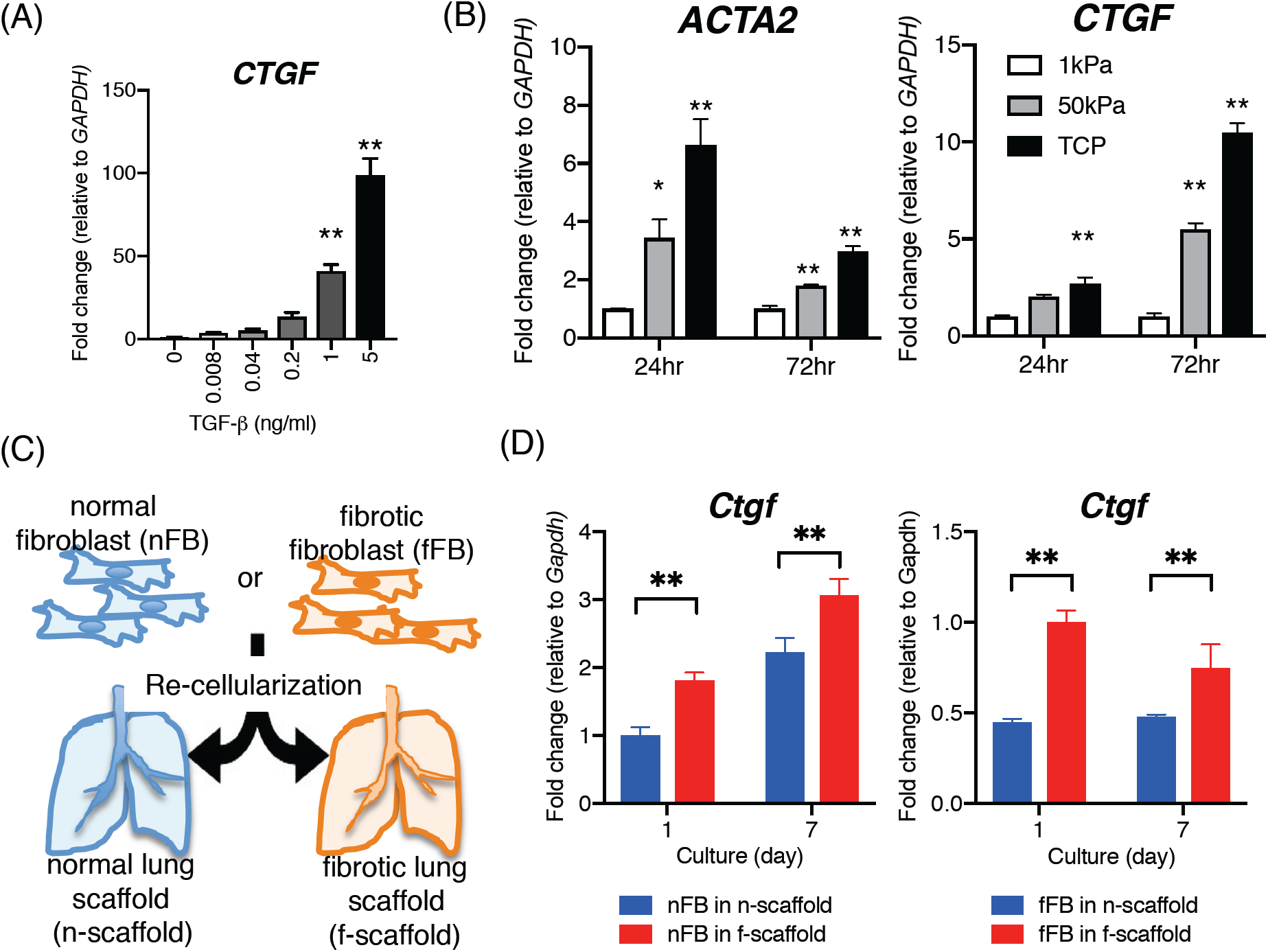
The regulatory mechanism of CTGF in the cells. (A) mRNA expression of *CTGF* in human lung fibroblasts treated with recombinant TGF-β for 24 hrs under serum free condition. (B) mRNA expression of *ACTA2* and *CTGF* in human lung fibroblasts cultured on different stiffness plates. (C) Schematic images of recellularization with rat fibroblasts in a rat decellularized lung and (D) mRNA expression of *Ctgf* in the recellularized lung at indicated time point. \*\**p* < 0.01, \**p* < 0.05.

The effect of matrix stiffness on CTGF expression in human lung fibroblasts was evaluated by culturing the cells on matricies of different stiffness (1 kPa, 50 kPa, and tissue culture plate [TCP]). The physiological stiffness of lungs ranges from 0.2–2 kPa, whereas pathological stiffness in the fibrotic lung ranges from 2–35 kPa (11)(12). Tissue culture plastic represents an extremely stiff matrix that is similar to tendon or bone, in the 10^6^ kPa range (13). A physiological stiffness is reported to keep lung fibroblasts in a quiescent state, whereas a pathological stiffness as in the fibrotic lungs is reported to induce a pro-fibrogenic phenotype with high proliferation and ECM synthesis rates (11). Within 24 hour of plating human lung fibroblasts on TCP, the expression of *ACTA2* mRNA was strongly elevated (Figure 4B). It was also elevated at 24 hours in cells on the 50 kPa plates, relative to cells plated on the soft matrix. By 72 hr, the expression of *ACTA2* mRNA on the stiffer matrices was lower than at 24 hours, but was still significantly elevated relative to that in cells on the soft matrix. The expression of *CTGF* mRNA was also induced in cells on the stiff plates, but with a somewhat different time course. At 24 hours after plating, *CTGF* mRNA was only significantly elevated on TCP. However, by 72 hr, *CTGF* mRNA was significantly greater in the fibroblasts plated on the 50 kPa matrix as well (5.5 times on 50 kPa and 10.5 times on TCP compared to on 1 kPa for 72 hrs) (Figure 4B). These data demonstrate that the stiffness surrounding the cells regulates CTGF expression.

To further investigate the effect of a 3D environment on CTGF expression, primary rat fibroblasts were exposed to normal or fibrotic lung scaffolds. Rat lungs were decellularized at 28 days after instillation of AdDL or AdTGF-β1 to obtain normal and fibrotic lung scaffolds, respectively. The lung scaffolds were then recellularized with primary rat fibroblasts isolated from normal and fibrotic lungs (Figure 4C). Post recellularization on day 1 and day 7, expression of *Ctgf* mRNA was examined. *Ctgf* mRNA expression was elevated in normal fibroblasts in a fibrotic scaffold, as compared to normal fibroblasts in a normal scaffold (Figure 4D). Interestingly, fibrotic fibroblasts in a normal scaffold exhibited reduced *Ctgf* mRNA expression (about 45% less) than that in fibrotic scaffolds at both time points. This observation indicates the plasticity of fibrotic fibroblasts responds to the microenvironment, and illustrates the importance of ECM in fibrogenesis (Figure 4D).

### Pamrevlumab attenuated TGF-β1-induced lung fibrosis in rats

In order to evaluate the importance of CTGF in lung fibrosis induced by TGF-β, AdTGF-β1 rats were therapeutically administered the anti-CTGF antibody, pamrevlumab, or placebo human antibodies. While Masson’s trichrome staining of lung sections on day 28 after instillation of AdTGF-β1 suggested that administration of pamrevlumab resulted in less lung remodeling and fibrosis, there was not a statistically significant reduction in the Ashcroft score on day 28 (Figure 5A). IF staining of lung sections showed that pamrevlumab administration reduced the expression of CTGF in Vimentin-positive cells (fibroblasts) (Figure 5B). Western Blot analyses were used to confirm the effect of pamrevlumab on CTGF expression and to evaluate its effect on the expression of other markers of fibrogenesis and TGF-β signaling: Collagen type 1, α-SMA, TGF-β, phospho-Smad3 (pSmad3), Smad3 and α-tubulin (Figure 5C). Pamrevlumab treatment resulted in a statistically significant reduction in the expression of CTGF, which is concordant with IF staining. Pamrevlumab also significantly decreased the abundance of α-SMA. There was a trend for reduced TGF-β in the lungs of pamrevlumab-treated rats, which was not statistically significant. However, canonical TGF-β signaling, as indicated by the pSmad3/Smad3 ratio, was significantly decreased in rats treated with pamrevlumab (Figure 5D).

**Figure 5.**
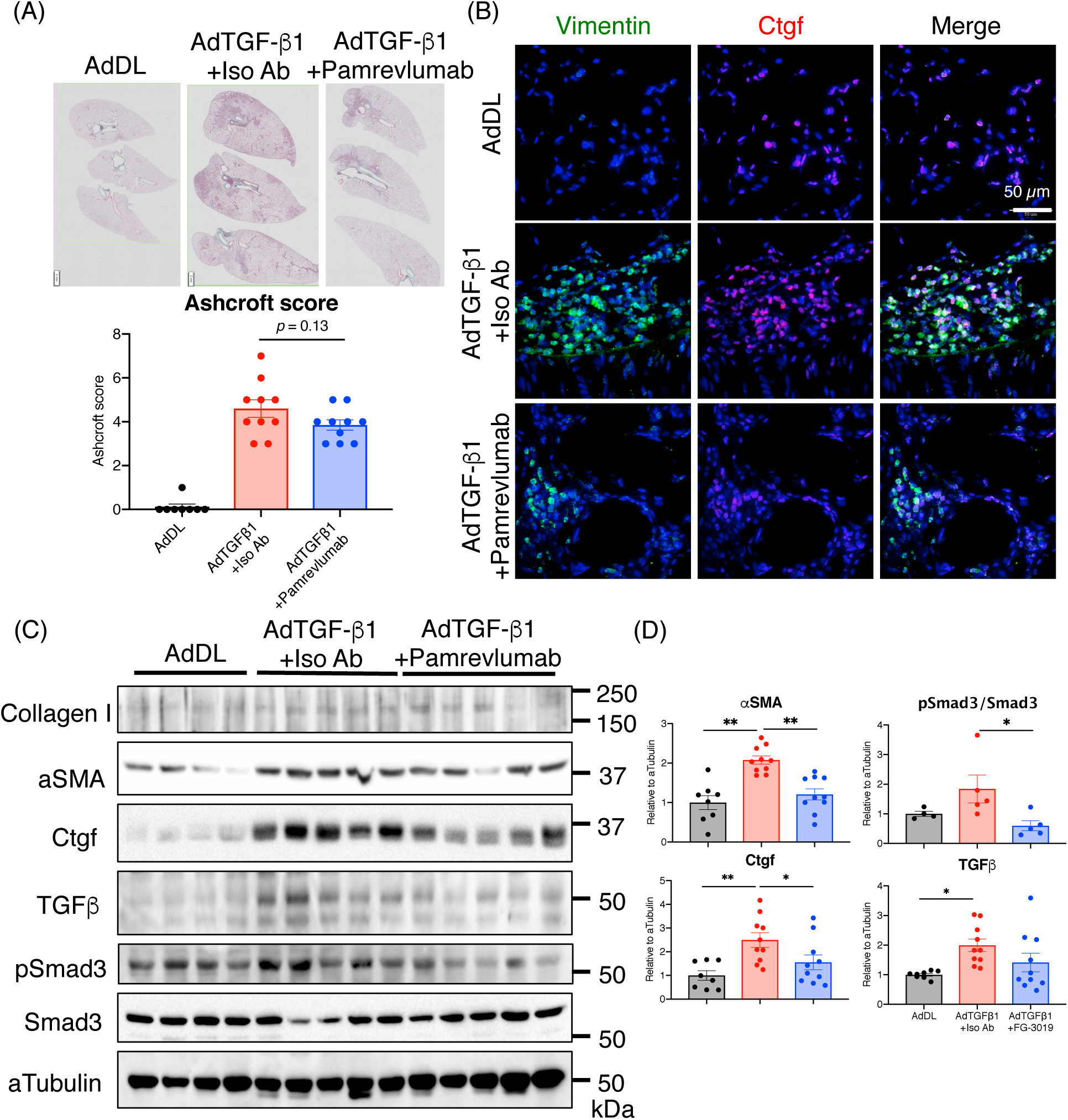
Pamrevlumab partially attenuated TGF-β1-induced lung fibrosis in rats. (A) Representative lung sections stained by Masson’s trichrome imaged by Slide Scanner and Ashcroft scoring from AdDL, AdTGF-β1+isotype control IgG (iso Ab), and AdTGF-β1+ pamrevlumab on day 28. (B) Immunofluorescence staining of Vimentin, CTGF, and DAPI on the lung sections from rats treated with AdDL, AdTGF-β1+iso Ab, and AdTGF-β1+ pamrevlumab on day 28. (C) Western blot analysis of Collagen type I, αSMA, CTGF, TGF-β, phospho-Smad3 (pSmad3), Smad3, and αTubulin in whole lung lysates of AdDL, AdTGF-β1+iso Ab, and AdTGF-β1+pamrevlumab on day 28. αTubulin was used as a loading control. AdDL; *n*=8, AdTGF-β1+iso Ab; *n*=10, AdTGF-β1+pamrevlumab: *n*=10. Data are expressed as mean ± SEM. \*\**p* < 0.01, \**p* < 0.05.

## Discussion

CTGF is a key mediator of tissue remodeling and fibrosis (14). CTGF activates myofibroblasts to produce pro-fibrotic extracellular matrix proteins leading to tissue remodeling and fibrosis (14). CTGF was previously reported to be elevated in the lungs of IPF patients and to be predominantly expressed in myofibroblasts and type 2 alveolar epithelial cells (15). In the present study, we observed a trend for elevated CTGF in lungs of IPF patients, which did not reach statistical significance due to variability. At the present time, it is unclear if the variability of CTGF expression observed in IPF lungs represents heterogeneity of the composition of the small pieces of lung examined, or whether it might depend on the epigenetic or genetic background of the patients, as previously reported in patients with systemic sclerosis (16). If CTGF expression in IPF patients is regulated by epigenetics or genetics, such information could assist with prescription of personalized therapy for treatment of IPF using an agent such as pamrevlumab. However, at the present time, there are no data to support this hypothesis.

Data from mining of single cell sequencing of normal and IPF lungs has greatly expanded the known number and types of cells in the lung that express CTGF. Many of these cells, such as fibroblasts, myofibroblasts, endothelial cells and smooth muscle cells, would have been predicted from cells known to express CTGF. Others, however, such as ciliated, club and goblet cells, appear to be novel. It is also interesting that CTGF expression in some of the highest expressing cells, such as myofibroblasts and venous endothelial cells, are very similar in healthy and IPF lungs. Only a few cell types exhibited substantial differences in CTGF expression between healthy and IPF lungs, including ciliated cells and fibroblasts, which both exhibited higher CTGF expression in IPF, and smooth muscle cells, which exhibited lower CTGF expression in IPF lung. How these changes relate to IPF pathophysiology remain to be elucidated.

CTGF was upregulated in multiple cell types with different time courses in the rat AdTGF-β1-induced lung fibrosis model. VSMC expressed elevated CTGF earlier than endothelial cells, which upregulated CTGF in the middle to later stages of fibrosis. Endothelial cell expression of CTGF is necessary for development of PH secondary to chronic hypoxia, and is a potential mechanism in the regulation of Rho family GTPase, Cdc42 (17). Our results suggest that altered vasculature cells actively contribute to pulmonary fibrosis through CTGF in cooperation with fibrotic fibroblasts. We speculate that interaction between vasculature cells and fibroblasts determine fibrotic features during the pathogenesis of IPF. Uncoordinated and uncontrolled release of CTGF by fibrotic cells may exaggerate abnormal tissue and vascular repair and subsequent formation of pulmonary fibrosis and pulmonary hypertension.

CTGF is an identified coordinator of vascular repair and vascular remodelling, which is an essential requirement for life as a complex organism (18). A variety of cells, from bone marrow-derived endothelial progenitor cells to peripheral circulating endothelial cells, serve as remote progenitors that can participate with locally proliferating cells in the formation of new blood vessels in response to a wide array of proangiogenic factors (18). Ischemia/hypoxia is a primary inducer of many of these factors, as are other stress and injury response pathways (18). CTGF is a hypoxia inducible protein (19)(20) with multiple cellular targets (18). Levels of plasma CTGF were also reported to be a potential biomarker in IPF, where CTGF levels were reported to be significantly elevated in IPF patients compared with non-IPF and healthy volunteers, and to correlate with loss of lung function (21).

Several studies have confirmed the cooperative effects of TGF-β and CTGF in fibrosis (22–26). CTGF has been reported to bind to TGF-β through the N-terminal von Willebrand factor domain, and this has been suggested to augment TGF-β activity, potentiating fibrosis (21). Our previous study, which showed that transient CTGF overexpression by itself is not sufficient to cause lung fibrosis, also strengthens the importance of the cooperative effects of TGF-β and CTGF (27).

Our data demonstrate the crucial role of the cellular environment for regulation of CTGF expression in 2D and 3D settings. Matrix stiffness influences the subcellular location of Transcriptional coactivator with PDZ-binding motif (TAZ) (25). Matrix stiffness facilitates F-actin organization, mechanotransduction, and activation of TAZ (25). TAZ in turn forms complexes with Smad2/3 that translocate into the nucleus and regulate target gene transcription following stimulation with TGF-β stimulation (25). These cellular responses could mediate the increased expression of CTGF and fibrotic markers in fibroblasts cultured on matrices of increased stiffness.

In the rat AdTGF-β1-induced lung fibrosis model, pamrevlumab treatment significantly decreased markers of fibrogenesis such as α-SMA and CTGF. Pamrevlumab also significantly decreased canonical TGF-β signaling, as indicated by reduced pSmad3. Since CTGF has been reported to directly affect non-canonical, but not canonical TGF-β signaling (28), the observation of pamrevlumab effects on canonical TGF-β signaling is likely the result of the decreased expression of TGF-β in this TGF-β-driven model. While visual inspection of Masson’s trichrome-stained sections suggested that pamrevlumab attenuated pulmonary remodeling and fibrosis, Ashcroft scoring was not able to confirm a statistically significant reduction of fibrosis. The rapid time course of this rodent model, and the short duration of treatment may have contributed to the limited response observed for pamrevlumab.

The results of a randomized, double-blind, placebo-controlled phase 2 clinical trial testing pamrevlumab in IPF patients (PRAISE) was recently reported (29). This trial, which was performed at 39 medical centres in seven countries (Australia, Bulgaria, Canada, India, New Zealand, South Africa and the USA), demonstrated that pamrevlumab reduced the decline in percentage predicted forced vital capacity by 60.3% at week 48 (mean change from baseline –2.9% with Pamrevlumab vs –7.2% with placebo) (29). The proportion of patients with disease progression was found to be lower in patients treated with pamrevlumab than in the placebo group at week 48 with good tolerability of pamrevlumab and a safety profile similar to that of placebo (29).

CTGF is upregulated in other fibrosing diseases/models: radiation-induced pulmonary fibrosis mouse model (30)(31), fibroblasts from systemic sclerosis patients (32)(33), bronchoalveolar lavage cells from patients with chronic sarcoidosis (34). A polymorphism in the CTGF promoter region has been reported to be associated with systemic sclerosis (16). Homozygosity for the G allele in the promoter of the *CTGF* gene (rs6918698) carries an increased risk of scleroderma and, in particular, an increased risk of interstitial lung fibrosis among patients with this disorder (16). These results suggest that upregulation of CTGF is a common feature in the pathogenesis of fibrosis, and therefore pamrevlumab could be an option for the treatment of pulmonary fibrosis.

## Acknowledgements

We thank Ms. Fuqin Duan for her technical assistance.

